# Codon arrangement modulates MHC-I peptides presentation: implications for a SARS-CoV-2 peptide-based vaccine

**DOI:** 10.1101/2021.02.04.429819

**Authors:** Tariq Daouda, Maude Dumont-Lagacé, Albert Feghaly, Alexandra-Chloé Villani

## Abstract

Among various vaccination strategies, peptide-based vaccines appear as excellent candidates because they are cheap to produce, are highly stable and harbor low toxicity. However, predicting which MHC-I Associated Peptide (MAP) will ultimately reach cell surface remains challenging, due to high false discovery rates. Previously, we demonstrated that synonymous codon arrangement (usage and placement) is predictive of, and modulates MAP presentation. Here, we apply CAMAP (Codon Arrangement MAP Predictor), the artificial neural network we used to unveil the role of codon arrangement in MAP presentation, to predict SARS-CoV MAPs. We report that experimentally identified SARS-CoV-1 and SARS-CoV-2 MAPs are associated with significantly higher CAMAP scores. Based on CAMAP scores and binding affinity, we identified 48 non-overlapping MAP candidates for a peptide-based vaccine, ensuring coverage for a high proportion of HLA haplotypes in the US population (>78%) and SARS-CoV-2 strains (detected in >98% of SARS-CoV-2 strains present in the GISAID database). Finally, we built an interactive web portal (https://www.epitopes.world) where researchers can freely explore CAMAP predictions for SARS-CoV-1/2 viruses. Collectively, we present an analysis framework that can be generalizable to empower the rapid identification of virus-specific MAPs, including in the context of an emergent virus, to help accelerate target identification for peptide-based vaccine designs that could be critical in safely attaining group immunity in the context of a global pandemic.

## Introduction

Since the emergence of SARS-CoV-2 in December 2019 in Wuhan city, Hubei province, China, scientific communities worldwide have engaged in a race against time to better understand the SARS-CoV-2 virus and develop anti-viral therapies and vaccines against COVID-19. The SARS-CoV-2 virus is a novel *Betacoronavirus* with high similarity to SARS-CoV-1^1^, another *Betacoronavirus* that caused a large disease outbreak of severe acute respiratory syndrome in 2002-2004.

Several studies have started to describe activation of T cell subsets in acute COVID19 patients^2–4^. Furthermore, virus-specific memory T cells, and especially cytotoxic CD8^+^ T cells, have been shown to play a pivotal role in both short- and long-term protection against SARS-CoV-1 infection^5,6^. To identify virus-infected cells, CD8^+^ T cells interact with MHC-I molecules, which present at the cell surface short peptides derived from the intracellular protein content. These MHC-I associated peptides (MAPs) represent important candidates for vaccine targets, as peptide-based vaccines present several advantages compared to other vaccine strategies: they contain no virulent factors, are very stable, can be easily produced and are capable of generating potent immune response^7^. Pre-clinical evidence have shown that peptide-based vaccines could provide immunity against SARS-CoV-1 infections, as Zhao and colleagues observed earlier virus clearance and increased survival in mice following immunization with SARS-CoV-1 peptide-pulsed dendritic cells^6^. However, a significant challenge with peptide-based vaccines is to obtain a combination of MAPs that will allow high coverage of the population, given that MHC-I alleles are highly polymorphic and that each allele binds a different peptide motif.

Identification of virus-specific MAPs remains difficult, especially in the context of an emergent virus. While predicting virus-specific MAPs can in theory be performed through analysis of viral genomic sequences, current predictive methodologies have shown very high false discovery rates^8–10^. Rules that regulate the binding of MAPs to MHC-I molecules have been well characterized using artificial neural networks (ANN) and weighted matrix approaches^11,12^. However, not all MAPs that can bind to an MHC-I allele will be naturally processed by the MHC-I antigen presentation machinery and ultimately reach the cell surface^13^. Indeed, while all proteins contain peptides that are predicted to bind MHC-I molecules, a recent study showed that as many as 41% of expressed protein-coding genes generated no MAPs^13^. Importantly, these authors also provided compelling evidence that the presentation of MAPs cannot be explained solely by their affinity to MHC-I alleles and their transcript expression levels, while ruling out low mass spectrometry sensitivity as an explanation for the non-presentation of the strong binders. Therefore, gaining a better understanding of the rules governing the biogenesis of naturally processed MAPs would prove extremely useful to determine which potential MAP would be useful vaccine targets, given that emergent viruses can be easily and rapidly sequenced.

Joining the facts that (1) MAPs appear to preferentially derive from defective ribosomal products (DRiPs)^14–19^ and (2) codon usage influences both precision and efficiency of protein translation^20,21^, we previously demonstrated that synonymous codon arrangement (usage and placement) is predictive of, and modulates MAP presentation^22^. We developed an artificial intelligence algorithm called Codon Arrangement MAP Predictor (CAMAP) that predicts MAP presentation from mRNA sequences flanking the MAP-coding codons (MCCs) and predicting MAP presentation in human cells with greater accuracy than when analyzing amino acid sequences. Most importantly, combining CAMAP scores with other known predictors of MAP presentation (i.e. MAP binding affinity to MHC-I molecules and/or transcript expression levels) significantly increased MAP prediction accuracy, and significantly decreases prediction false positive rate^22^.

Here, we sought to determine whether the rules that CAMAP derived on human mRNA sequences to predict naturally processed MAPs also apply to viral sequences. As SARS-CoV-2 shows high genomic similarity to the SARS-CoV-1 virus^1^, we elected to analyze both SARS-CoV-1/2 mRNA sequences and extract CAMAP scores for all their potential MAPs. We first confirm that naturally processed SARS-CoV-1 specific MAPs (identified since the pandemic of 2002-2004), and mass spectrometry identified SARS-CoV-2 MAPs^23^ are indeed associated with high CAMAP scores. We then used both CAMAP scores and predicted MHC-I binding affinity to define a set of 15 very common HLA class I alleles in order to identify a combination of 48 high potential SARS-CoV-2-specific MAPs. These 48 peptides were characterized by high CAMAP scores and strong binding affinities to several frequent HLA alleles. Analysis of the predicted binding affinity of these 48 selected MAPs to the 15 most common HLA alleles showed that all haplotypes containing one of these common HLA alleles were predicted to present at least one selected MAP in the US population, with a median of 14 MAPs being presented by each haplotype. Interestingly, a third of our selected MAPs were homologous to SARS-CoV-1 naturally processed peptides previously identified, suggesting that our selection method could accelerate target identification for peptide-based vaccines. Finally, in an effort to facilitate data sharing and exploration, we created an interactive web platform called www.epitopes.world on which the complete list of MAPs from both SARS-CoV-1/2 viruses and their associated CAMAP scores can be explored.

## Results

### CAMAP framework used to predict naturally processed MAPs

CAMAP predicts MAP presentation from flanking MAP-coding codons (MCCs). The algorithm receives as input mRNA sequences and computes probabilities that these sequences yield naturally processed MAPs. CAMAP was developed using a previously published dataset consisting of MAPs presented on B-lymphoblastic cell line (B-LCL) isolated from 18 subjects and expressing a total of 33 MHC-I alleles^13,24^. Following their detection by mass spectrometry, MAPs were associated to their source transcripts using pyGeno^25^ to generate a hit and decoy datasets, which were used for training and validation of the CAMAP algorithm^22^. Noteworthy, CAMAP scores are independent from, and complementary to MAP MHC-I binding affinity^22^. Therefore, the list of potential MAPs predicted by CAMAP should then be filtered according to their capacity to bind the specific MHC-I alleles expressed by individuals to create a list of MAPs with high potential, which are defined as MAPs that have a high likelihood of being naturally processed and presented at the cell surface by the specific MHC-I molecules. Figure 1 recapitulates the CAMAP algorithm and analyses frameworks that we used here to predict SARS-CoV-1/2 MAPs.

**Figure 1.**
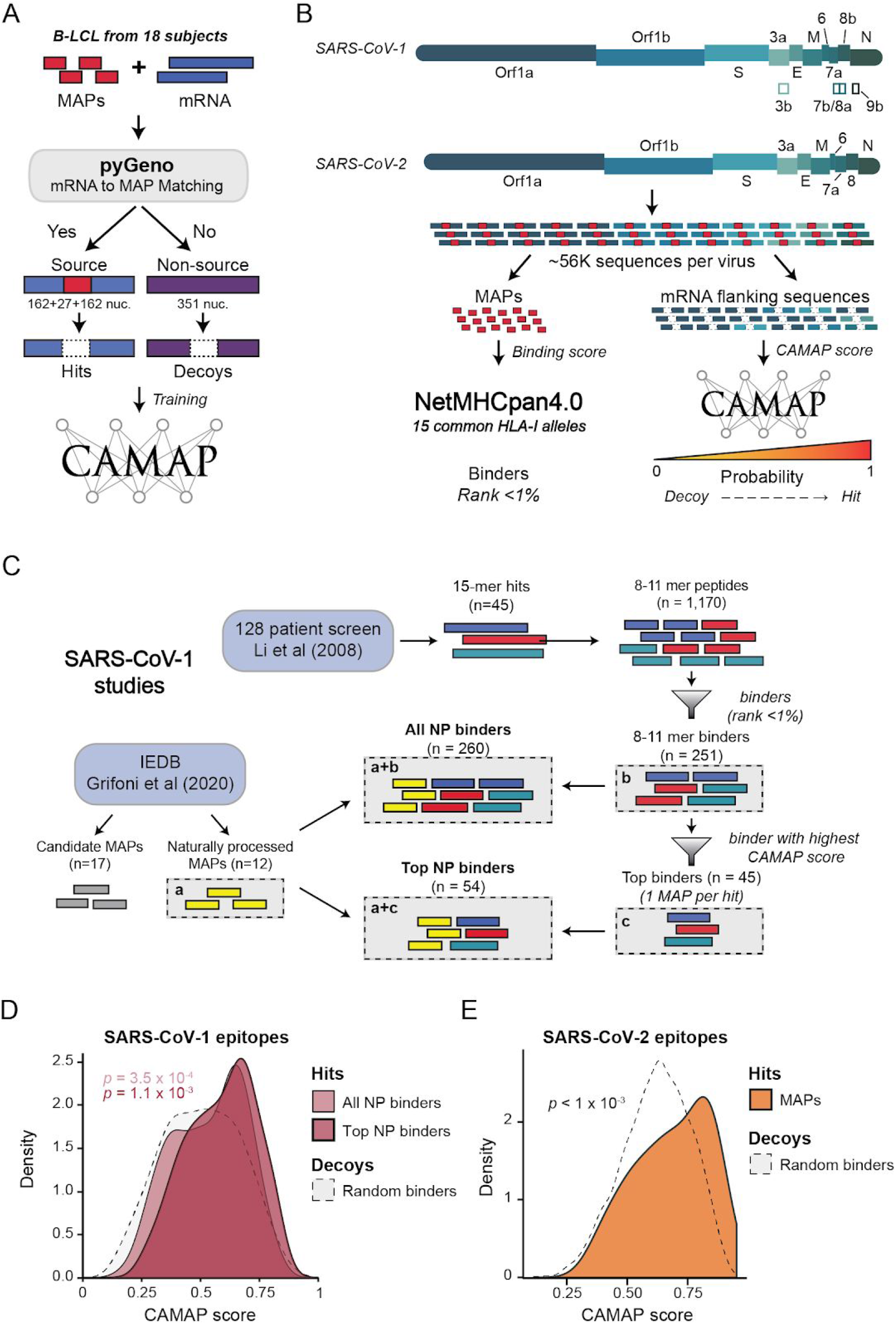
SARS-CoV-1/2 naturally processed MAPs show high CAMAP scores compared to random SARS-CoV-1/2 binders (i.e. minimal ranks < 1%). (A) CAMAP algorithm training (see Methods for more details). (B) Evaluation of CAMAP score and MAP binding affinity (NetMHCpan4.0) for SARS-CoV-1/2 coding sequences and potential MAPs. The CAMAP score represents the probability that an mRNA sequence gives rise to a MAP, with a probability of 1 indicating this sequence is very likely to generate a MAP. (C) Dataset description of candidates and naturally processed (NP) SARS-CoV-1 MAPs. The first subset of SARS-CoV-1 MAPs was compiled from the IEDB database and contains 17 candidates and (**a**) 12 naturally processed SARS-CoV-1 MAPs^1^. The second subset derived from the study of CD8^+^T cell response in 128 patients who recovered from SARS-CoV-1 infection^28^. These authors screened the whole SARS-CoV-1 genome using 15-mer peptides. From this screen, they identified 45 hits which induced a specific CD8^+^T cell response in ≥1 patient in their cohort. As MHC-I molecules bind shorter peptides of 8-11 amino acids, we derived all linear 8 to 11-mer peptides from these 15-mer hits (n=1,170), which were then filtered for binders (**b**, n=251), i.e. with a minimal rank <1% for ≥1 common HLA alleles (Supplementary Table S1). Then, for each 15-mer hit, we selected the binder (from **b**) with the highest CAMAP score (one binder per hit only), for a total of 45 top binders (**c**). We referred to the combination of **a** and **b** as “All naturally processed (NP) binders”, and of **a** and **c** as “Top naturally processed (NP) binders”. Three MAPs were found in all subsets combined. (D-E) SARS-CoV-1^1,28^ (D) and SARS-CoV-2^23^ (E) naturally processed (NP) MAPs are associated with higher CAMAP scores than decoy binders (i.e. MAPs with minimal binding score rank <1%) originating from the same proteins. We defined the calculated *p*-value as the proportion of times where the median of the decoy dataset was either equal or superior to that of the hit dataset (one-tailed).

To evaluate the likelihood that the MAP presentation rules derived by CAMAP on human sequences also apply to viral sequences, we compared CAMAP predictions for all possible SARS-CoV-1/2 MAPs to the predictions given by CAMAPs on its reserved test set^22^. We extracted all possible MAPs (lengths of 8-11 amino acids) and their associated mRNA flanking sequences (we used CAMAP’s default context size of 162 nucleotides on each side) for both SARS-CoV-1 and SARS-CoV-2 viruses (NCBI: NC_004718 and NC_045512). We then computed the CAMAP score associated with the mRNA flanking sequence of each MAP (Fig. 1A-B). In addition, we used NetMHCpan4.0^26^ to predict MHC-I binding affinity of each MAP to the 15 most common HLA class I alleles (HLA-A, -B and -C) in the US population^27^ (Supplementary Table S1) to identify MAPs most likely to be presented by MHC-I molecules.

### Naturally processed SARS-CoV MAPs have high CAMAP scores

It has previously been shown that integrating CAMAP score into MAP prediction frameworks significantly reduced the false positive rate in predicting human MAPs^22^. Here, we evaluate whether this also translates to a more accurate prediction of naturally processed SARS-CoV-1/2 MAPs. We first compiled a list of 71 unique SARS-CoV-1 MAPs previously described in the literature (Fig. 1C and Supplementary Table S2). We separated them into two categories: (i) naturally processed MAPs (n=54) (i.e. MAPs that have been demonstrated to be naturally processed and presented at the surface of SARS-CoV-1 infected cells), and (ii) candidate MAPs (n=17) (i.e. MAPs that are known to bind at least one MHC-I allele but for which we could not find evidence of the candidate being naturally processed). Because no proteogenomic analysis of SARS-CoV-1 infected cells has been reported hitherto, we used the detection of MAP-specific CD8^+^ T cells response in patients who recovered from a SARS-CoV-1 infection as a proxy for the detection of naturally processed MAP.

The first list of SARS-CoV-1 MAPs was recently compiled from the IEDB database by Grifoni *et al* (2020), and contains both candidates (n=17) and naturally processed MAPs (n=12, see Fig. 1C)^1^. The second list of MAP hits (n=45) was described by Li et al (2008) and derived from the study of CD8^+^T cell response in 128 recovering patients from SARS-CoV-1 infection using *ex vivo* IFN-gamma ELISPOT assays. The authors used an unbiased approach in which they screened the whole SARS-CoV-1 genome using a matrix of 15-mers overlapping peptides^28^. However, as MHC-I molecules bind shorter peptides of 8 to 11 amino acids, it is generally assumed that these 15-mer peptides will undergo additional processing before being presented at the cell surface^29^. Thus, we derived all possible 8- to 11-mer MAPs from each 15-mer hit and filtered them for binders, which we define as MAPs predicted to bind at least one common HLA allele with a minimal rank <1% (n=251, see Fig. 1C). In addition, for each 15-mer hit, we extracted the binder with the highest CAMAP score, which we refer to this set of MAPs as “top binders” (Fig. 1C). Thus, we generated two lists of naturally processed SARS-CoV-1 MAPs by combining the 12 naturally processed MAPs compiled by Grifoni et al (2020) with either all naturally processed binders or top naturally processed binders derived from the 15-mer hits described by Li et al (2008) (Fig. 1C). Of note, three MAPs were identified in both studies.

We then compared the distribution of CAMAP scores between these two sets of candidates (All and Top naturally processed binders) with those of random binders derived from the same proteins (Fig. 1D). Both sets of candidates showed significantly higher CAMAP scores than random binders, demonstrating that the rules derived by CAMAP also apply to viral sequences. Finally, top naturally processed MAPs are associated with statistically significant higher CAMAP scores than other candidate MAPs (Supplementary Figure S1). Indeed, 72% (39/54) of top naturally processed MAPs showed a CAMAP score greater than the median CAMAP score of their respective protein of origin, compared to only 41% of candidate MAPs (*p* = 0.039, Fisher exact test).

More recently, Weingarten-Gabbay et al. (2020) described 30 MAPs specific to SARS-CoV-2 in A549 (lung) and HEK293 (kidney) SARS-CoV-2 infected cell lines^23^. Of these 30 MAPs, 27 originated from canonical open-reading frames (ORFs), and 3 were from internal out-of-frame ORF in the Spike and Nucleoprotein genes. In addition, 6 of them originated from ORF9b, one of the 23 novel recently identified ORFs^30^. SARS-CoV-2 specific MAPs showed high CAMAP scores compare to random binders derived from the same protein origin (*p* < 1 x 10^−3^, Figure 1E). Indeed, 15/21 (71%) of them showed a CAMAP score that was above the median CAMAP scores of their respective protein (Supplementary Figure S2), similar to what was observed with SARS-CoV-1-specific MAPs. Taken together, these results support the use of CAMAP scores in predictions of MAPs from viral genetic sequences.

### Replicase and Spike proteins as likely sources of MAPs

SARS-CoV-1/2 sequences both showed very distinct CAMAP score distributions when compared to our baseline of human sequences used to train CAMAP. Indeed, while the majority of the human B-LCL-derived sequences (the decoys of the dataset on which CAMAP was trained) have both low CAMAP scores and low affinity to MHC-I alleles (in gray in Fig. 2A), most SARS-CoV-1/2 viral sequences showed high CAMAP scores (i.e. >0.5), resembling the distribution of hits in the training datasets (i.e., sequences associated with naturally processed human MAPs), but have low MAP binding affinity (Fig. 2A). Noteworthy, when feeding CAMAP with shuffled sequences (i.e. mRNA sequences where codons are randomly replaced with synonymous codons to erase any codon specific information^22^), predictions for both viruses become centered around the neutral value 0.5. This result shows that CAMAP scores derived from SARS-CoV sequences is strongly influenced by codon arrangement, contrarily to naturally processed human sequences for which shuffled sequences still retain some predictive power^22^. As expected, due to their very high homology^1^, distribution of CAMAP scores and binding affinities were almost identical for both SARS-CoV viruses (Fig. 2A).

**Figure 2.**
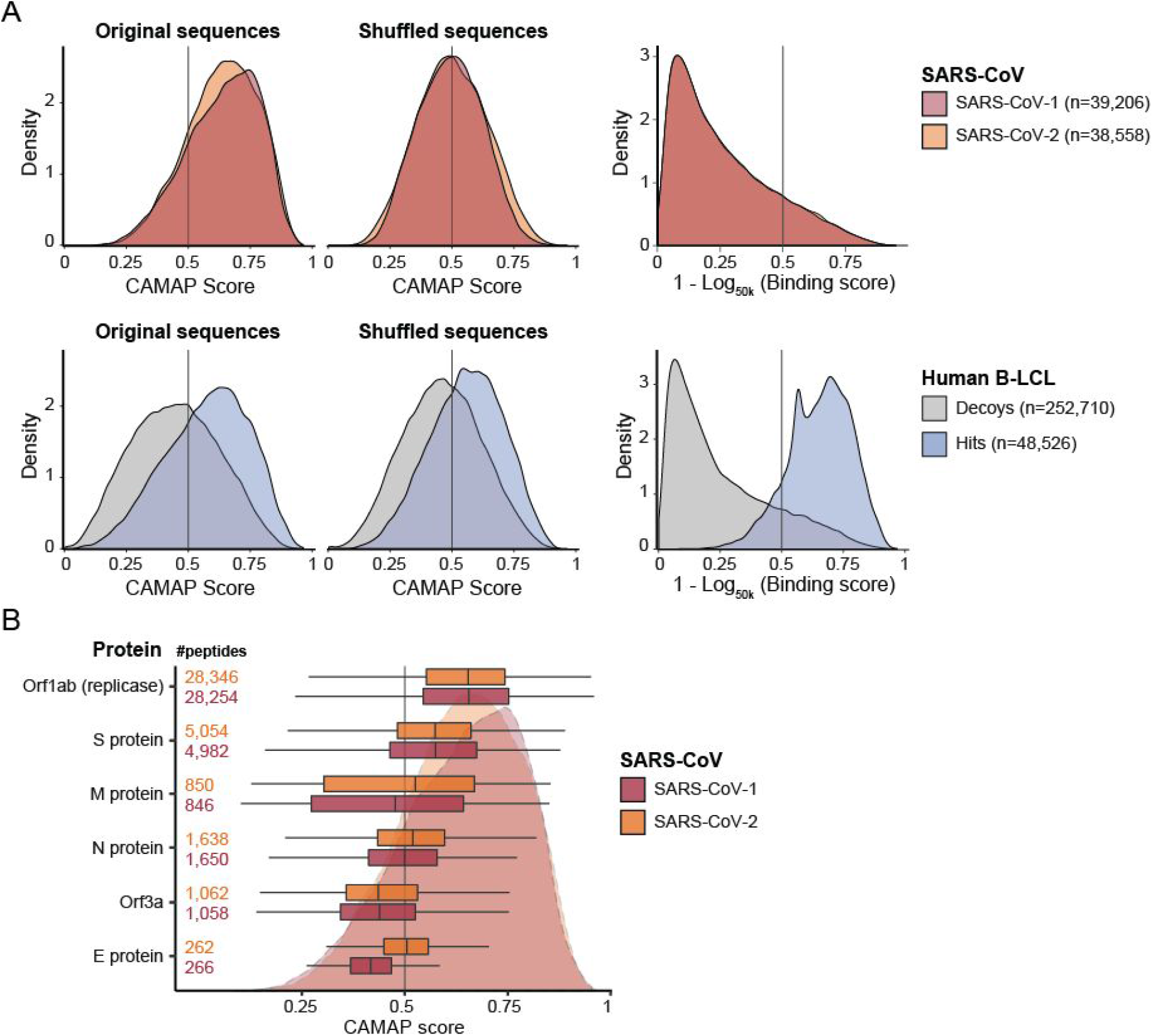
Replicase and S proteins of both SARS-CoV-1/2 have high CAMAP scores, suggesting that they could be preferential sources of MAPs. (**A**) Distribution of CAMAP scores (left panels) and minimal peptide binding affinity (i.e. the minimal binding score predicted across 15 common alleles right panel) for SARS-CoV-1/2 (upper panels) and Hit and Decoy sequences from B-LCL (bottom panels). Here, a low binding score means low affinity for MHC-I molecules, while a binding score close to one means high affinity (transformed using 1 – log_50,000_(BS) where BS is expressed in nM). (**B**) CAMAP score distributions for the 5 main SARS-CoV-1/2 proteins.

The Replicase (Orf1ab), which occupies two thirds of SARS-CoV-1/2 genomes, and the Spike protein coding sequences both had particularly high CAMAP score distributions compared to the other viral genes (Fig. 2B). These results suggest that SARS-CoV-1/2 viruses contain numerous sequences that have the potential to encode for naturally processed MAPs, with most of them being derived from the Replicase (Orf1ab) and Spike protein coding sequences.

### Design of a SARS-CoV-2 peptide vaccine

We next designed a strategy to rank-order potential SARS-CoV-2 MAPs according to their likelihood of being naturally presented in the greatest proportion of possible individuals. Our framework involved first focusing our selection criteria on binders (i.e. peptides with a minimal rank <1% for at least one HLA allele) that were derived from the 5 main proteins encoded in the SARS-CoV-2 genome: Orf1ab (replicase complex), Spike (or S) protein, matrix (M) protein, envelop (E) protein and the nucleocapsid (N) protein. We first filtered peptides with a CAMAP score above the 75^th^ percentile compared to other peptides derived from the same protein. Of note, as the E protein is very short (75 amino acids) and thus contains a low number of potential MAPs (only 53 binders), we lowered our filter threshold to the 50^th^ percentile of CAMAP scores for this protein. To select MAPs that can be presented in a high proportion of individuals, we calculated the cumulative frequency of haplotypes (HLA-A, -B and -C) capable of presenting each given MAP. The haplotypes that are capable of presenting a given peptide were defined as containing at least one HLA gene with a predicted binding score below 1% (rank, NetMHCpan4.0^26^). We focused only on the 15 most common HLA alleles in the US population (Supplementary Table S1), which captured approximately 78.9% of haplotypes described for the US population^27^. Through this strategy, MAPs predicted to bind very frequent HLA alleles (e.g. HLA-A*02:01) and/or several HLA alleles were thus prioritized in our ranking. Of note, when several binders overlapped one another with >75% identity, we selected the binder predicted to be presented by the highest cumulative frequency of haplotypes.

Following these selection criteria, we report herein the top 10 MAPs for each of the 5 main SARS-CoV-2 proteins, which we believe are particularly appealing for peptide-based vaccine targets. The 48 high potential SARS-CoV-2 MAPs have high CAMAP scores and are predicted to be presented in a high proportion of individuals (E protein: n=8, other proteins: n=10 each, see Table 1). Of note, 14 out of 48 (29.2%) of our selected SARS-CoV-2 MAPs are homologous (with an overlap >50%) to peptides that were previously identified as naturally processed SARS-CoV-1 MAPs^1,28^ (indicated with a star in Table 1). Furthermore, 3 out of 48 peptides that we identified as SARS-CoV-2 targets MAPs (see Table 1) had >65% identity (i.e. overlapping amino acids) by mass spectrometry of SARS-CoV-2 infected cells^23^ (Supplementary Table S3). In contrast, when MAPs are rank-ordered only on the basis of their capacity of being presented in a high proportion of individuals (i.e. rank-ordering only on the basis of binding affinities, without pre-filtering for high CAMAP scores), only 10/48 (20.8%) MAPs correspond to homologues of SARS-CoV-1 naturally processed MAPs (Supplementary Table S4). These results suggest that including CAMAP scores in MAP predictions enriches for naturally processed peptides.

**Table 1.**
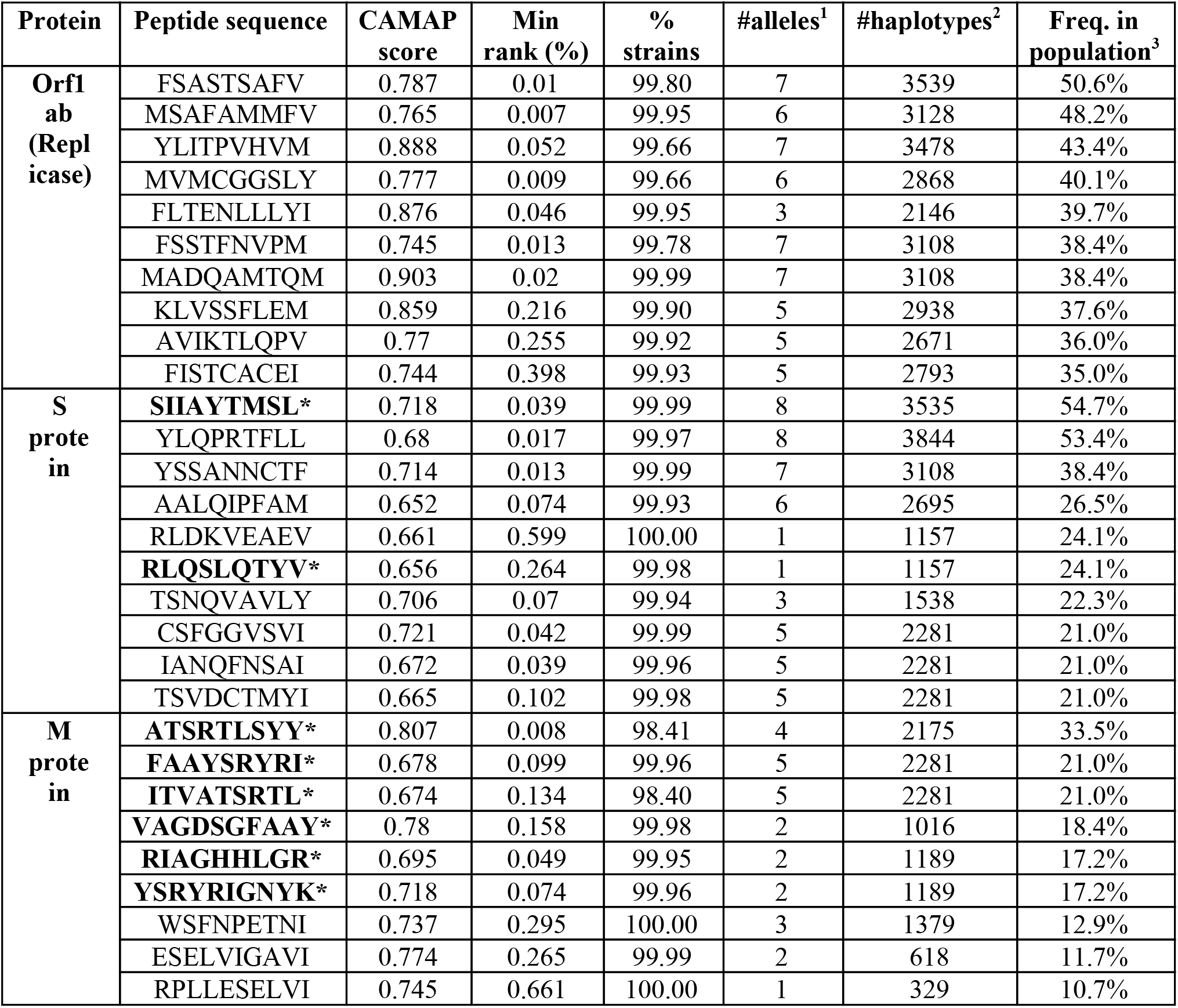
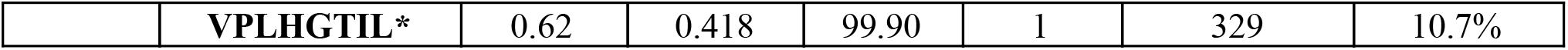
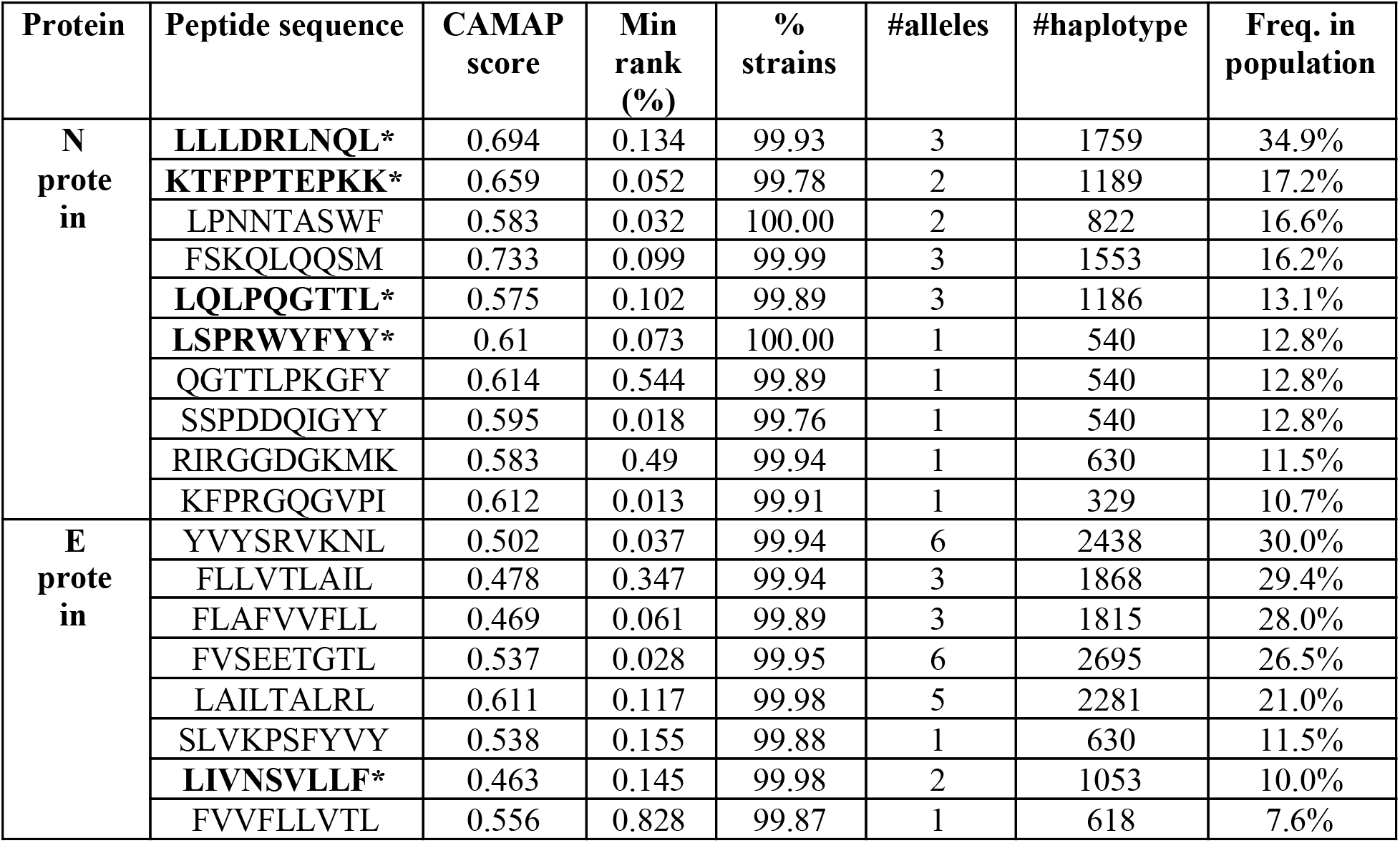
List of top SARS-CoV-2 target MAPs. For each peptide sequence (organized by SARS-CoV-2 proteins), we listed: its associated CAMAP score, its minimal rank (binding affinity) to 15 most common HLA alleles, the proportion of different strains that contains each MAP sequenced as of June 8^th^ 2020 (as extracted from GISAID database), the number of common HLA alleles that it can bind to with a rank <1% (1), and the number of haplotypes that can present it with at least one allele in the US population (2). Last column shows the overall frequency in the US population of haplotypes predicted to bind each peptide with at least one common HLA allele (3). These peptides were further verified as having an absence of shared sequence homology with the human or murine proteins using the BLAST search engine http://www.ncbi.nlm.nih.gov/blast/, to prevent any autoimmune response. 14/48 (29.2%) of target peptides were homologous (or with an overlap >50%) to those that were previously identified as naturally processed SARS-CoV-1 MAPs^1,28^ (indicated with a star: *).

Most of our high CAMAP score selected MAPs (37/48, 77%) are predicted to bind multiple HLA alleles (Table 1, Fig. 3A, Supplementary Table S5). These selected MAPs have strong binding affinities (low ranks) and high CAMAP scores (Fig. 3B and C). Haplotypes containing one of the 15 common HLA alleles are predicted to present a median of 14 MAPs (range 1-35, Fig. 3D), with >75% of haplotypes presenting at least 5 MAPs. Our selected MAPs: (i) have a high potential of being naturally processed, (ii) are predicted to be presented by very common HLA class I molecules, (iii) have been detected in more than 98% of strains sequenced across the world since the beginning of the pandemic (until June 8, 2020, data downloaded from the GISAID database; Table 1) and (iv) exhibit little to no variation in their CAMAP scores caused by missense or silent mutations in their surrounding contexts of 162 nucleotides as observed in those strains (Supplementary Figure S3). These results suggest that filtering for MAPs with high CAMAP scores could accelerate identification of naturally processed MAPs originating from viral sequences. If immunogenic, these MAPs could be used as targets for a peptide-based vaccine against SARS-CoV-2, that would be compatible with a high proportion of the population.

**Figure 3.**
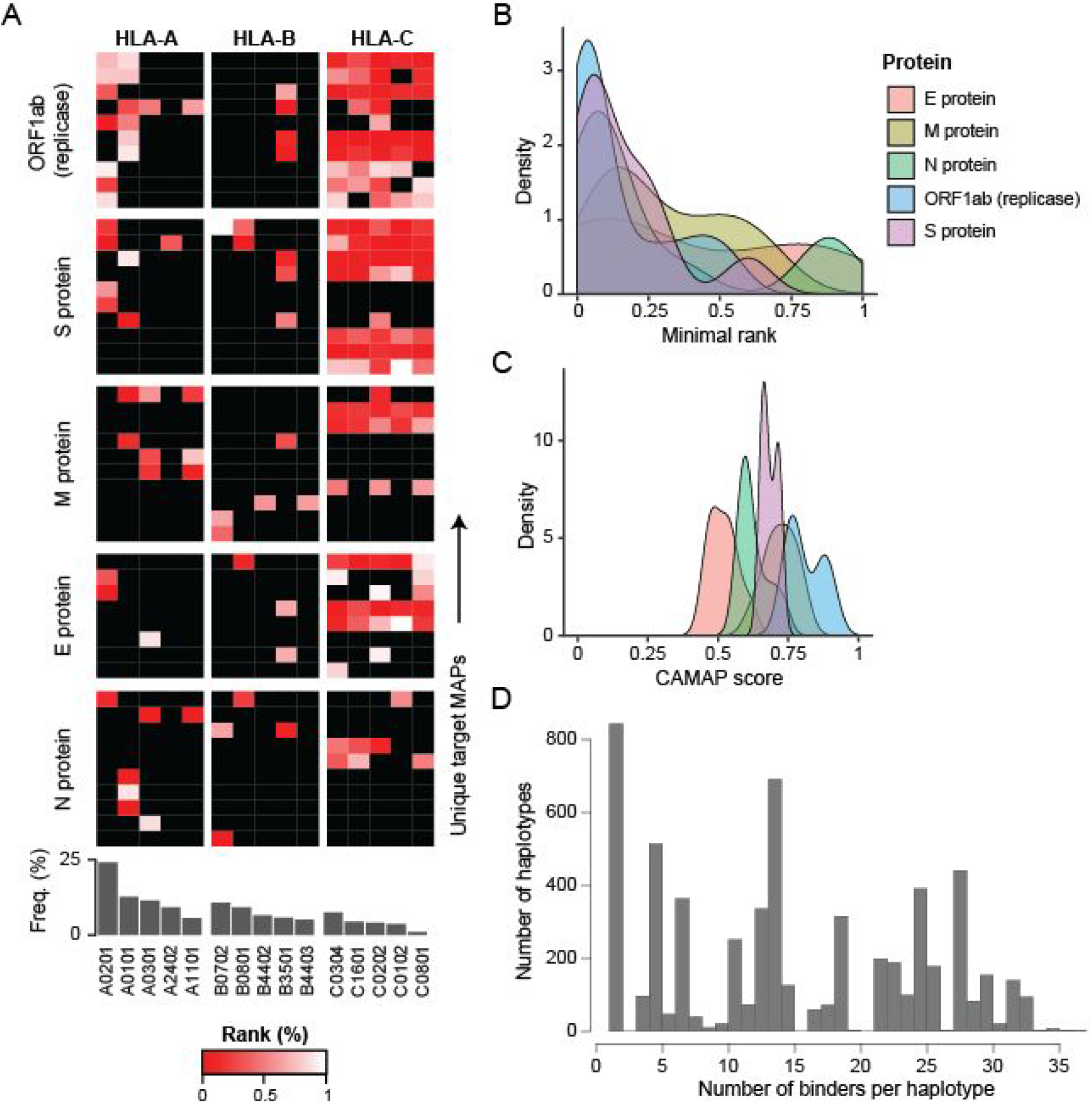
Presentation of selected SARS-CoV-2 MAPs (n=48) across frequent haplotypes in the US population. (**A**) MAPs binding affinity to specific HLA-A, -B and -C alleles (red to white gradient). Black indicates that MAPs have a rank ≥1% for this specific allele. Alleles are ordered according to their frequency in the US population (shown in the bar graph below). (**B-C**) Distribution of minimal binding scores (rank) (**B**) and CAMAP scores (**C**) for selected MAPs from each of the five main SARS-CoV-2 proteins. (**D**) Number of selected MAPs predicted to be presented by unique haplotypes (i.e. MAPs with a minimal rank <1% for at least one HLA class I allele in haplotype).

### Interactive web platform

To foster collaboration and in an effort to speed up the discovery of a peptide-based SARS-CoV-2 vaccine, we have built an interactive web platform allowing researchers to freely explore and export CAMAP results on SARS-CoV-1/2 (Fig. 4A-B). Our platform is hosted on https://www.epitopes.world and contains all predictions made on the genomes of SARS-CoV-1, SARS-CoV-2 (about 39,000 peptides each, Fig. 4B). For reference, we have also included predictions made on B-LCLs (the original dataset CAMAP was trained and tested on), for both hit and decoy sequences (a total of about 104,000 peptides). The portal hosts a total of over 180,000 unique peptides and 379,000 entries (many human peptides have several variable mRNA flanking regions). Through epitopes.world, researchers can filter for organism, protein, peptide length, and CAMAP score, plot distributions of CAMAP scores and export the corresponding lists of peptides with their CAMAP score in a csv file. The interface also allows overlaying several CAMAP score distributions for a user-friendly experience.

**Figure 4.**
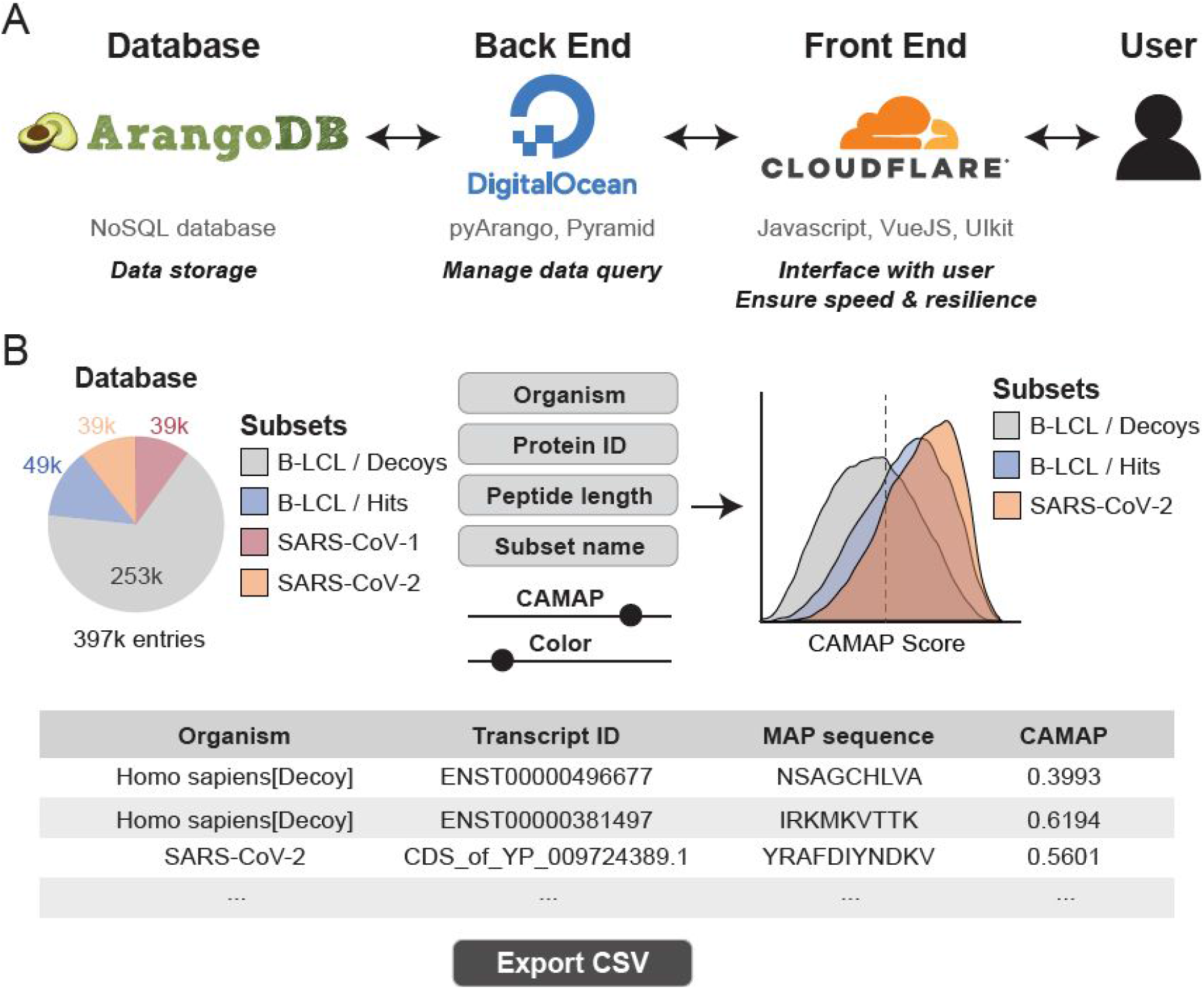
Interactive web portal epitopes.world. (**A**) To ensure scalability and high availability of the results we designed a custom cloud infrastructure: (1) All data is permanently stored in an instance of ArangoDB running on the ArangoOasis cloud solution; (2) query results are served through a python implemented REST API hosted on the cloud provider DigitalOcean; (3) We used the Content Delivery Network (CDN) Cloudflare to serve the JavaScript front-end that provides the graphical user interface, and to cache the results of frequent queries. (**B**) Upon selecting which dataset to plot (defined by organism, protein ID and/or peptide length), peptides can be filtered for their CAMAP score and length. Users can then define a plot name and choose a color for the selected dataset. Multiple datasets can be plotted simultaneously and the resulting peptide list is displayed in a table below.

## Discussion

Predicting naturally processed MAPs from the analysis of genetic sequences has historically resulted in very high rates of false positive^8–10^. In a recent study, Daouda *et al*. (2020) showed that the mRNA sequences flanking the MAP-coding codons were predictive of MAP presentation, and complementary to binding affinity and mRNA expression, supporting the role of co-translational events in the regulation of MAP biogenesis. In this study, we show that combining predicted binding affinities and CAMAP scores increases the prediction accuracy of virus-derived naturally processed MAPs, which we define as MAPs that are naturally processed by the antigen presentation machinery and are therefore presented by MHC-I molecules at the surface of virus infected cells. Indeed, we showed that naturally processed MAPs previously detected on SARS-CoV-1 infected cells are associated with higher CAMAP scores compared to other potential binders derived from the same proteins (i.e. MAPs with a minimal binding score, or rank, <1%).

We also showed that mass-spectrometry identified SARS-CoV-2 MAPs were also associated with higher CAMAP scores. Of note, in Weingarten-Gabbay et al. study, the 2 proteins that gave rise to the highest number of MAPs were ORF1ab (11/30, 37%) and the S proteins (7/30, 23%), while only 1 peptide originated from each of the M and N proteins^23^. This result is very surprising, as the mRNA expression levels of the ORF1a and ORF1ab is 10 to 1000-fold lower than that of structural ORFs^30^. However, this is in accordance with the general distribution of CAMAP scores (see Figure 2B), which are highest for ORF1ab, followed by the S protein.

Based on these findings, we designed a framework to rank-order potential MAPs according to their CAMAP scores (representing their likelihood of being naturally processed by the antigen presentation machinery) and their capacity to be presented across several very common HLA alleles. We also confirmed that each nominated MAP through our selection criteria had been detected in more than 98% of the ~40,000 strains sequenced and available in the GISAID database on June 8, 2020. We report herein the top 10 MAPs for each of the main SARS-CoV-2 proteins (only 8 MAPs for the E protein, due to its small size), which we believe are particularly appealing for peptide-based vaccine targets. Interestingly, ~1/3 (29.2%) of these SARS-CoV-2 selected MAPs were homologous to naturally processed peptides previously identified on SARS-CoV-1 infected cells^1,28^. In contrast, when rank-ordering SARS-CoV-2 MAPs only based on their capacity to be presented across several common HLA alleles, only ~1/5 (20.8%) of the selected MAPs were homologous to SARS-CoV-1 naturally processed MAPs. These results suggest that filtering for MAPs with a high CAMAP score could significantly accelerate the identification of naturally processed MAPs, which would be a highly useful feature for the rapid development of peptide-based vaccines. Further tests will be required to determine the immunogenicity of each of these peptides and to validate whether these peptides can indeed be detected at the surface of infected cells.

In the context of a global pandemic, vaccines remain the only means of safely attaining group immunity. When new viruses emerge, as is the case with SARS-CoV-2, vaccine design needs to be optimized for four main parameters: (1) minimal toxicity, (2) high population coverage, (3) rapid development and validation, and (4) efficient production and distribution. Our approach presents several advantages regarding these four parameters. First, the low toxicity of peptide-based vaccines makes them attractive thanks to their high safety profiles. Secondly, our method of epitope selection guarantees that a high proportion of the population can benefit from the vaccine, which is necessary to achieve group immunity, and that each selected MAP has been found in more than 98% of strains sequenced since the beginning of the pandemic. Thirdly, using CAMAP scores should increase the rate of true positive predictions, thus decreasing costs and accelerating vaccine development. Finally, peptide-based vaccines can be synthetized quickly and relatively cheaply, ensuring a rapid production of the vaccine, and are highly stable^7^, facilitating worldwide distribution. Although we have here limited ourselves to the analysis of SARS-CoV-1/2 viruses, our results suggest that similar analysis framework to the one we have presented for the identification and prioritization of candidate MAPs can be implemented and generalized in the future to other viruses. Theses could be critical in helping accelerating target identification for peptide-based vaccine designs in the context of a global pandemic.

## Methods

### CAMAP training

CAMAP algorithm was trained using mRNA sequences flanking MAP-coding codons (MCCs). MAPs derived from B-LCL from 18 subjects were detected by mass spectrometry and paired to their mRNA sequences of origin using pyGeno. These sequences constituted the Hit dataset. Decoy dataset was derived from mRNA sequences that did not encode any MAPs. For both the Hit and Decoy datasets, the MCCs were removed from each mRNA sequence, leaving only the MAP-flanking codons (with a context size of 162 nucleotides).

### SARS-CoV-1/2 sequences

Genetic coding sequences of SARS-CoV-1/2 viruses were extracted from NCBI (NC_004718.3 and NC_045512.2), and parsed using the BioPython library^31^. SARS-CoV-1/2 unique mRNA contexts (MAP-flanking) and their associated MAP were extracted. MAP binding affinity to 15 common HLA class I alleles were predicted using NetMHCpan4.0, while CAMAP score associated with each flanking sequence was extracted using CAMAP.

### Analysis of SARS-CoV-1 specific peptides

Epitopes associated with specific CD8^+^T cell responses were extracted from Li et al (2008) and Grifoni et al (2020)^1,28^. Epitopes were further classified either into (i) naturally processed (i.e. peptides for which there is evidence of natural processing and presentation), or (ii) potential candidates (i.e. peptides for which there is evidence of binding to MHC-I molecules and generation of specific cytotoxic CD8^+^ T cells only). Of note, peptides derived from synthetic sequences were categorized as potential candidates, as the mRNA sequences flanking the MAP-coding codons were different from the viral sequence.

Furthermore, Li et al (2008) used 15-mer peptides spanning the whole SARS-CoV proteome in their screening assay. However, MHC-I molecules bind peptides of 8 to 11 amino acids in length. Therefore, we extracted all possible linear 8 to 11-mer “sub-peptides” that can be derived from each 15-mers epitopes for which CD8^+^ T cell response was detected (n=44)^28^. We systematically extracted their corresponding binding score (NetMHCpan4.0^26^) and CAMAP score associated with their coding transcript. As the binding of MAPs to MHC-I molecules is necessary for MAP presentation, we kept only those sub-peptides that could bind (rank <1%) to at least one common HLA allele (frequency in the population >5%, Supplementary Table S1), as described by Maiers et al (2007)^27^. These 15 common HLA alleles are found in 78.9% of haplotypes described in the general US population.

All naturally processed (NP) binders and Top NP binders in Fig. 2A were defined as follow:

- All NP binders: these are defined as all the naturally processed peptides described in Grifoni *et al.* (2020) (n = 12) and all binders (minimal rank <1%) derived from Hits (15-mer epitopes) identified by Li *et al.* (2008) that can bind one of the 15 common HLA alleles (n = 276, Fig. 1C);
- Top NP binders: these are defined as all the naturally processed peptides described in Grifoni *et al.* (2020) (n = 12), and one binder (minimal rank <1%) per Hit identified by Li *et al.* (2008) with the highest CAMAP score among all binders derived from the same 15-mer Hit (n = 41, Fig. 1C).

As the different SARS-CoV proteins have different distributions of CAMAP scores and different lengths, we compared SARS-CoV-1 naturally processed binders to a set of random decoy peptides that (1) originated from the SARS-CoV-1 proteins in the same ratio than the hit datasets, and (2) were predicted to bind at least one common HLA allele (n=276). Of note, epitopes captured in the hit datasets were not removed from the decoy dataset. We then compared the median CAMAP score in the Hit and Decoy datasets using a bootstrap approach (n = 1,000,000), where the decoy dataset was resampled in each iteration. We defined the calculated *p*-value as the proportion of times where the median of the decoy dataset was either equal or superior to that of the hit dataset (one-tailed). The resulting fraction was reported as *p*-value in Fig. 1D-E.

### Selection of high potential SARS-CoV-2 MAPs

The description of haplotypes for the US African American, Asian, European American, and Hispanic populations was extracted from https://bioinformatics.bethematchclinical.org/hla-resources/haplotype-frequencies/^27^. The 15 most frequent HLA-A, -B and -C alleles (>5%) in the US population were defined in this study as the most common HLA alleles (Supplementary Table S1). SARS-CoV-2 MAPs’ binding affinity (rank) was extracted for those 15 alleles using NetMHCpan4.0^26^. Of those, we kept only peptides that had a minimal binding affinity <1% for at least one allele in our subset of common HLA-A, -B or -C alleles.

As each protein presented different distributions of CAMAP score, we filtered our candidate peptides to keep those with the highest CAMAP scores compared to other peptides within the same protein (i.e. top 25^th^ percentile for Orf1ab, S, M and N proteins, top 50^th^ percentile for E protein). We then computed the overall frequency of haplotypes that contain at least one common HLA allele (A, B or C) capable of binding a given peptide (rank <1%). We removed overlapping peptides (i.e. with >75% overlapping amino acids) to keep only the one with the highest overall frequency of binding haplotypes or the highest CAMAP score. We selected as high potential MAPs the top 10 peptides from each of the main SARS-CoV-2 proteins (i.e. ORF1ab replicase and structural proteins: S, M, E and N). As the E protein is very small, only 8 MAPs corresponded to our selection criteria. In consequence, we presented a list of 48 selected SARS-CoV-2 MAPs.

In addition, we confirmed that our selected MAPs were detected in more than 98% of all strains sequenced up until June 8, 2020 and available in the GISAID database (https://www.gisaid.org/). These consisted of 42,109 strains that were aligned to the reference SARS-CoV-2 genome from NCBI (NC_045512.2) using MUSCLE v3.8.31 with default parameters and the BioPython library. All peptide positions are reported in relation to NC_045512.

## Author Contributions

TD, AF and MDL contributed to data analysis, study design and performed computational experiments. TD and MDL co-wrote the first draft of the paper. ACV contributed to study design, reviewed and contributed to the manuscript. All co-authors reviewed the manuscript.

## Acknowledgements

We would like to thank members of the team who built the interactive web platform https://www.epitopes.world: Olivier Caron-Lizotte, Antoine Zieger, Logan Schwartz, Walter Sobral and Jörg Schad. We also would like to thank Barbara Decelle for her support and Explorai, ArangoDB and Digital Ocean for supporting the platform epitopes.world.

The authors declare no competing interests.

## Notes

### Competing Interest Statement

The authors have declared no competing interest.

## References

1. Grifoni, A. et al. A Sequence Homology and Bioinformatic Approach Can Predict Candidate Targets for Immune Responses to SARS-CoV-2. Cell Host Microbe 27, 671–680.e2 (2020).

2. Le Bert, N. et al. SARS-CoV-2-specific T cell immunity in cases of COVID-19 and SARS, and uninfected controls. Nature 584, 457–462 (2020).

3. Grifoni, A. et al. Targets of T Cell Responses to SARS-CoV-2 Coronavirus in Humans with COVID-19 Disease and Unexposed Individuals. Cell 181, 1489–1501.e15 (2020).

4. Weiskopf, D. et al. Phenotype and kinetics of SARS-CoV-2-specific T cells in COVID-19 patients with acute respiratory distress syndrome. Sci. Immunol. 5, (2020).

5. Channappanavar, R., Fett, C., Zhao, J., Meyerholz, D. K. & Perlman, S. Virus-specific memory CD8 T cells provide substantial protection from lethal severe acute respiratory syndrome coronavirus infection. J. Virol. 88, 11034–11044 (2014).

6. Zhao, J., Zhao, J. & Perlman, S. T cell responses are required for protection from clinical disease and for virus clearance in severe acute respiratory syndrome coronavirus-infected mice. J. Virol. 84, 9318–9325 (2010).

7. Li, W., Joshi, M. D., Singhania, S., Ramsey, K. H. & Murthy, A. K. Peptide Vaccine: Progress and Challenges. Vaccines 2, 515–536 (2014).

8. Nielsen, M., Lundegaard, C., Lund, O. & Keşmir, C. The role of the proteasome in generating cytotoxic T-cell epitopes: insights obtained from improved predictions of proteasomal cleavage. Immunogenetics 57, 33–41 (2005).

9. Abelin, J. G. et al. Mass Spectrometry Profiling of HLA-Associated Peptidomes in Mono-allelic Cells Enables More Accurate Epitope Prediction. Immunity 46, 315–326 (2017).

10. Capietto, A.-H., Jhunjhunwala, S. & Delamarre, L. Characterizing neoantigens for personalized cancer immunotherapy. Curr. Opin. Immunol. 46, 58–65 (2017).

11. Bassani-Sternberg, M. & Gfeller, D. Unsupervised HLA Peptidome Deconvolution Improves Ligand Prediction Accuracy and Predicts Cooperative Effects in Peptide–HLA Interactions. J. Immunol. 197, 2492–2499 (2016).

12. Nielsen, M. & Andreatta, M. NetMHCpan-3.0; improved prediction of binding to MHC class I molecules integrating information from multiple receptor and peptide length datasets. Genome Med. 8, (2016).

13. Pearson, H. et al. MHC class I–associated peptides derive from selective regions of the human genome. J. Clin. Invest. 126, 4690–4701 (2016).

14. Antón, L. C. & Yewdell, J. W. Translating DRiPs: MHC class I immunosurveillance of pathogens and tumors. J. Leukoc. Biol. 95, 551–562 (2014).

15. Wei, J. et al. Ribosomal Proteins Regulate MHC Class I Peptide Generation for Immunosurveillance. Mol. Cell 73, 1162–1173.e5 (2019).

16. Yewdell, J. W., Antón, L. C. & Bennink, J. R. Defective ribosomal products (DRiPs): a major source of antigenic peptides for MHC class I molecules? J. Immunol. 157, 1823–1826 (1996).

17. Croft, N. P. et al. Kinetics of antigen expression and epitope presentation during virus infection. PLoS Pathog. 9, e1003129 (2013).

18. Milner, E., Barnea, E., Beer, I. & Admon, A. The turnover kinetics of major histocompatibility complex peptides of human cancer cells. Mol. Cell. Proteomics MCP 5, 357–365 (2006).

19. Hassan, C. et al. The human leukocyte antigen-presented ligandome of B lymphocytes. Mol. Cell. Proteomics MCP 12, 1829–1843 (2013).

20. Cannarozzi, G. et al. A Role for Codon Order in Translation Dynamics. Cell 141, 355–367 (2010).

21. Plotkin, J. B. & Kudla, G. Synonymous but not the same: the causes and consequences of codon bias. Nat. Rev. Genet. 12, 32–42 (2011).

22. Daouda, T. et al. Codon arrangement modulates MHC-I peptides presentation. bioRxiv 2020.06.03.078824 (2020) doi:10.1101/2020.06.03.078824.

23. Weingarten-Gabbay, S. et al. SARS-CoV-2 infected cells present HLA-I peptides from canonical and out-of-frame ORFs. BioRxiv Prepr. Serv. Biol. (2020) doi:10.1101/2020.10.02.324145.

24. Granados, D. P. et al. Proteogenomic-based discovery of minor histocompatibility antigens with suitable features for immunotherapy of hematologic cancers. Leukemia 30, 1344–1354 (2016).

25. Daouda, T., Perreault, C. & Lemieux, S. pyGeno: A Python package for precision medicine and proteogenomics. F1000Research 5, 381 (2016).

26. Jurtz, V. et al. NetMHCpan-4.0: Improved Peptide-MHC Class I Interaction Predictions Integrating Eluted Ligand and Peptide Binding Affinity Data. J. Immunol. Baltim. Md 1950 199, 3360–3368 (2017).

27. Maiers, M., Gragert, L. & Klitz, W. High-resolution HLA alleles and haplotypes in the United States population. Hum. Immunol. 68, 779–788 (2007).

28. Li, C. K. et al. T cell responses to whole SARS coronavirus in humans. J. Immunol. Baltim. Md 1950 181, 5490–5500 (2008).

29. Zhang, H. et al. Comparing Pooled Peptides with Intact Protein for Accessing Cross-presentation Pathways for Protective CD8+ and CD4+ T Cells. J. Biol. Chem. 284, 9184–9191 (2009).

30. Finkel, Y. et al. The coding capacity of SARS-CoV-2. Nature (2020) doi:10.1038/s41586-020-2739-1.

31. Cock, P. J. A. et al. Biopython: freely available Python tools for computational molecular biology and bioinformatics. Bioinformatics 25, 1422–1423 (2009).

