## Supplemental materials for "Codon arrangement modulates MHC-I peptides presentation: implications for a SARS-CoV-2 peptide-based vaccine"

### Supplementary Figures

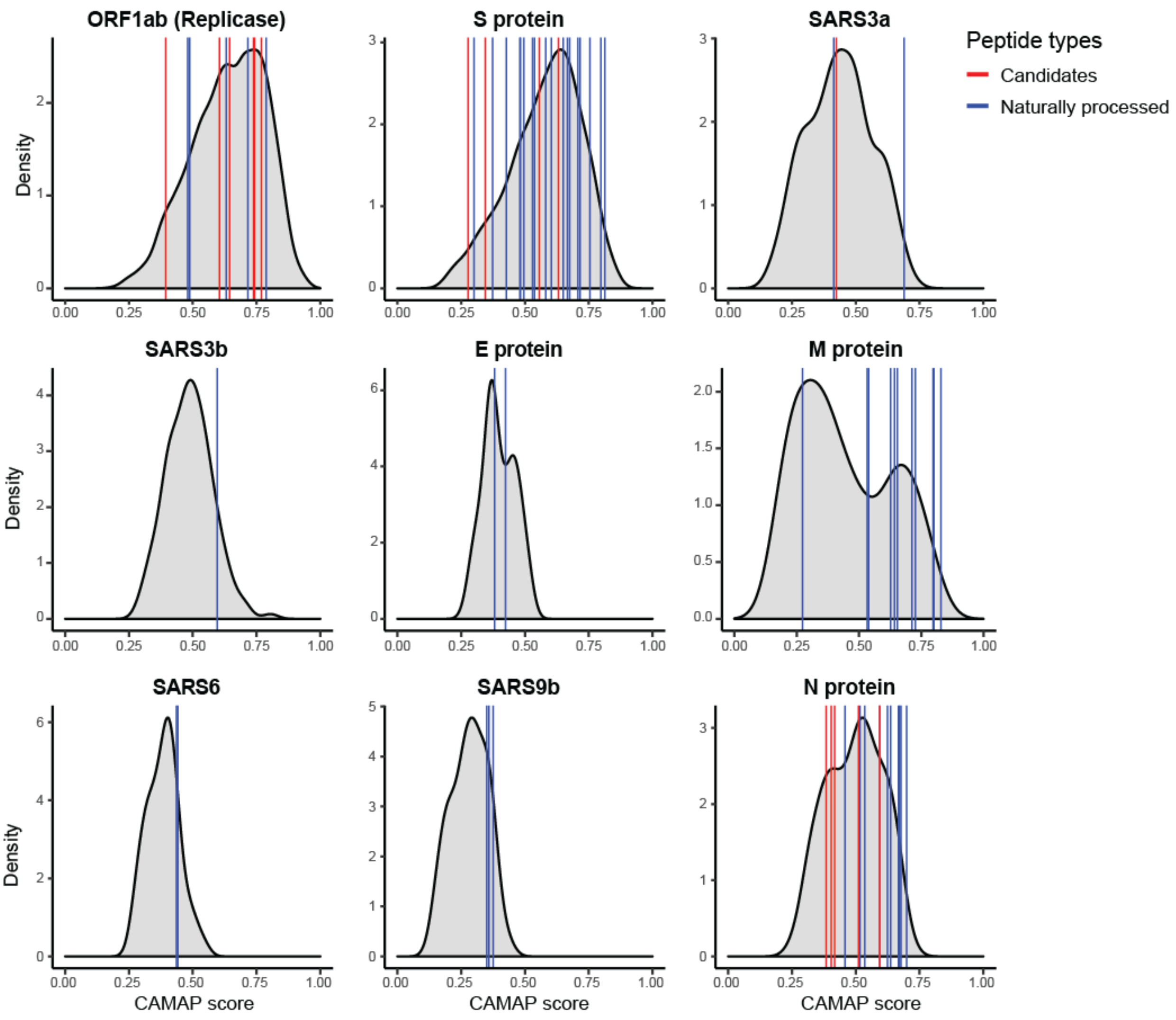


**Supplementary Figure S1. CAMAP score of top MAPs associated with SARS-CoV-1.** Only potential binders (peptides with a minimal rank <1% across 43 common HLA alleles) are shown for both Hits and global protein. Hits were separated into naturally processed (i.e. peptides for which there are evidence of natural processing and presentation from viral sequence) shown in blue, versus potential candidates (i.e. peptides for which there are evidence of binding to MHC-I molecules and generation of specific cytotoxic T cells only) shown in red. Of note, peptides which were derived from synthetic sequences were categorized as potential candidates, as the context flanking the MAP-coding codons were different from the viral sequence. Most naturally processed peptides have a CAMAP score above 0.5 or above the median CAMAP score for their protein of origin. Only the top binders derived from the study of Li *et al.* (2008) are presented here (see Fig. 1C).

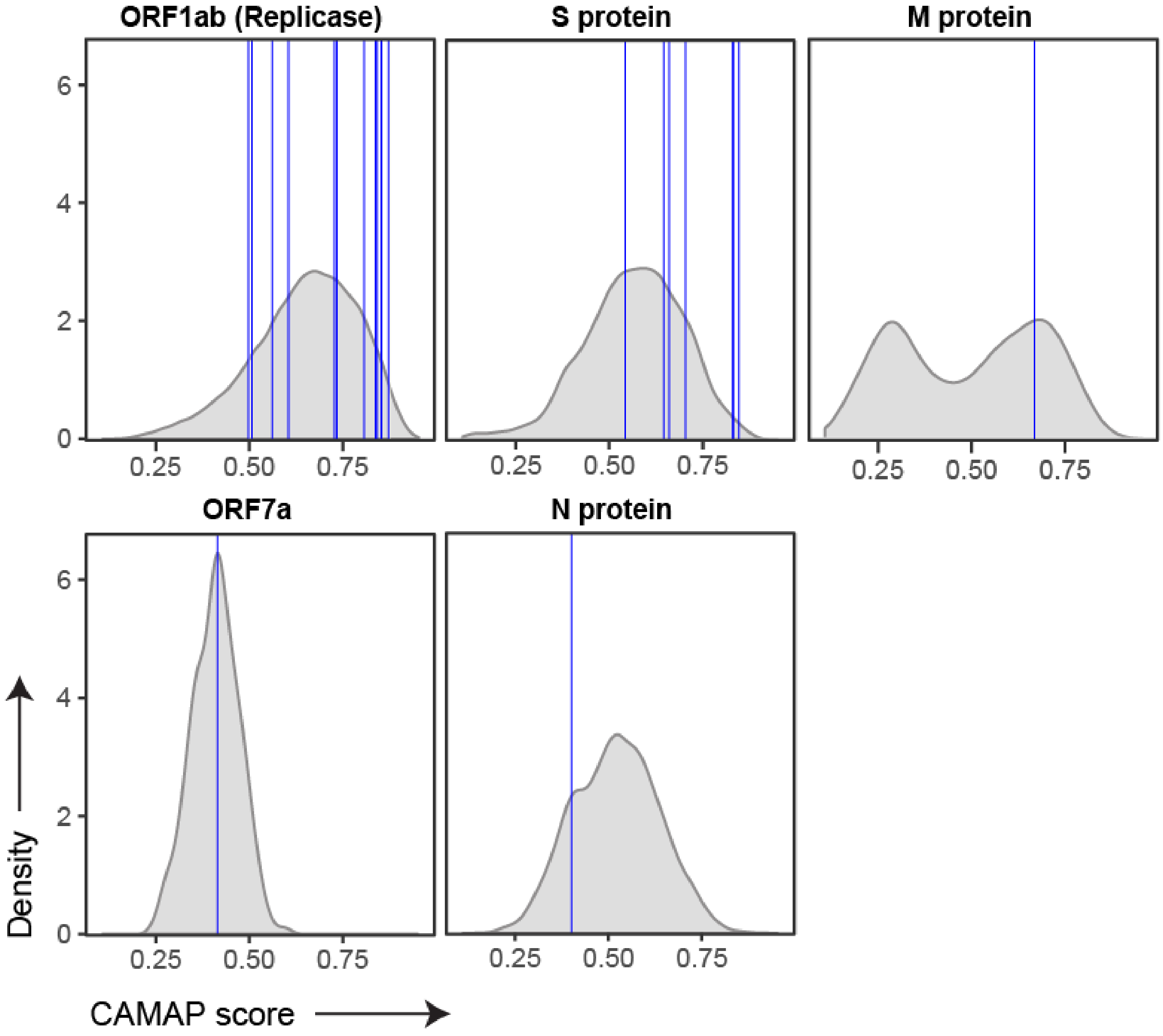


**Supplementary Figure S2. CAMAP score of MAPs presented by SARS-CoV-2 infected cells.** CAMAP scores of each SARS-CoV-2 specific naturally processed MAPs identified by mass spectrometry (Weingarten-Gabbay, S. et al, 2020) is shown with a blue line, and compared to the distribution of CAMAP scores of all potential binders (i.e. MAPs with a minimal binding score >1% rank) derived from the same protein.

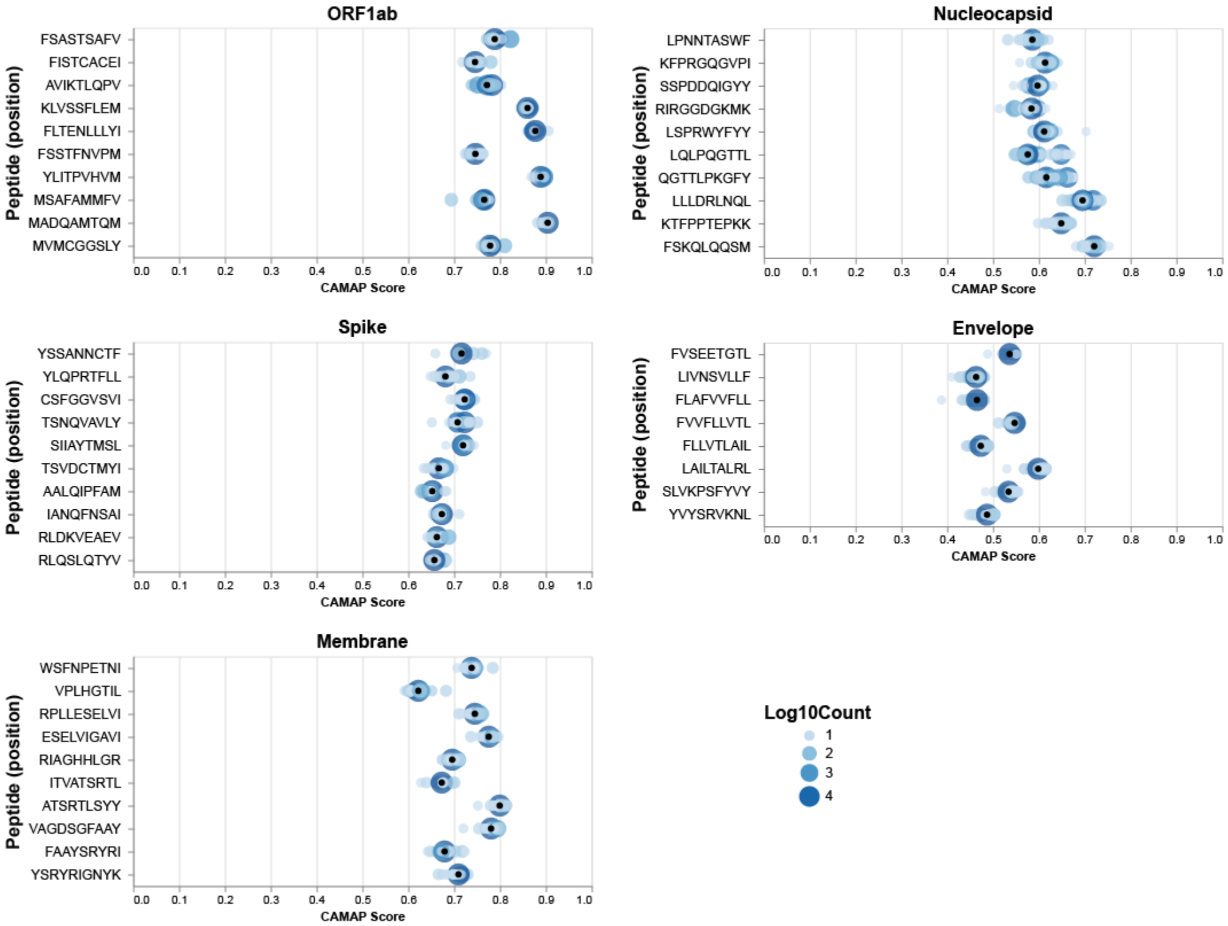


**Supplementary Figure S3. CAMAP score distributions for SARS-CoV-2 selected peptides.** All 48 selected peptides from the 5 main SARS-CoV-2 proteins (see Table 1) are included. Score distributions are from the ~40,000 GISAID strains used in this study. Black points represent the reference CAMAP score from NC_045512 that is used in the rest of this paper. Due to the extremely high number of data points, a log10 transformation was applied to the counts prior to plotting.

### Supplementary Tables

#### Supplementary Table S1. Most common HLA-A, -B and -C alleles in the US population.

| HLA | ALLELE | EUROPEAN | AFR.AMER | ASIAN/PAC.ILS. | HISPANICS | GLOBAL |
|---|---|---|---|---|---|---|
| A | **02:01** | 29.57% | 12.50% | 9.58% | 19.30% | **24.10%** |
| | **01:01** | 17.17% | 4.76% | 5.10% | 6.69% | **12.76%** |
| | **03:01** | 14.31% | 8.15% | 2.56% | 7.93% | **11.53%** |
| | **24:02** | 8.65% | 2.22% | 18.30% | 12.27% | **9.14%** |
| | **11:01** | 5.63% | 1.59% | 17.93% | 4.56% | **5.69%** |
| B | **07:02** | 13.97% | 7.26% | 2.69% | 5.51% | **10.74%** |
| | **08:01** | 12.51% | 3.90% | 1.63% | 4.51% | **9.15%** |
| | **44:02** | 8.99% | 2.14% | 0.77% | 3.28% | **6.47%** |
| | **35:01** | 5.69% | 6.52% | 4.25% | 6.24% | **5.81%** |
| | **44:03** | 4.95% | 5.41% | 4.24% | 6.11% | **5.19%** |
| C | **08:01** | 16.6% | 12.4% | 3.9% | 10.4% | **14.0%** |
| | **01:02** | 15.0% | 7.0% | 14.6% | 11.3% | **13.2%** |
| | **02:02** | 10.5% | 18.5% | 8.0% | 16.4% | **12.5%** |
| | **16:01** | 9.3% | 8.8% | 6.6% | 5.9% | **8.4%** |
| | **03:04** | 9.1% | 3.5% | 0.9% | 4.6% | **7.0%** |

**Supplementary Table S2. SARS-CoV-1 hits from [1] Li *et al.* (2008) and [2] Grifoni *et al.* (2020).** Of note, only top binders are shown in this table for MAPs derived from the study by Li *et al.* (2008).

| **Protein** | **SARS-CoV-1** | | | | **SARS-CoV-2** | | | **Identity (%)** | **Ref.** |
|---|---|---|---|---|---|---|---|---|---|
| | **Peptide** | **Category[1]** | **CAMAP score** | **Min. rank** | **Peptide** | **CAMAP score** | **Min. rank** | | |
| ORF1ab | WLMWFIISI | PC | 0.39 | 0.12 | WLMWLIINL | 0.46 | 0.18 | 67 | 2 |
| | CGYLPTNAV | NP | 0.48 | 0.26 | CGYLPQNAV | 0.62 | 0.39 | 91 | 1 |
| | SDGTGTIY | NP | 0.49 | 0.82 | SDGTGTIY | 0.68 | 0.82 | 100 | 1 |
| | ALSGVFCGV | PC | 0.60 | 0.05 | SLPGVFCGV | 0.48 | 0.12 | 78 | 2 |
| | IFVDGVPFV | NP | 0.63 | 0.30 | IFVDGVPFV | 0.61 | 0.30 | 100 | 1 |
| | SMWALVISV | PC | 0.64 | 0.03 | SMWALIISV | 0.56 | 0.02 | 89 | 2 |
| | ILLLDQALV | PC | 0.70 | 0.31 | ILLLDQALV | 0.76 | 0.31 | 100 | 2 |
| | TLKEILVTY | NP | 0.72 | 0.24 | TLKEILVTY | 0.61 | 0.24 | 100 | 1 |
| | CLDAGINYV | PC | 0.74 | 0.08 | CLEASFNYL | 0.60 | 0.47 | 56 | 2 |
| | TLMNVITLV | PC | 0.74 | 0.02 | TLMNVLTLV | 0.63 | 0.02 | 89 | 2 |
| | LLCVLAALV | PC | 0.77 | 0.99 | SACVLAAEC | 0.62 | 15.38 | 56 | 2 |
| | LLATNNVFRL | NP | 0.79 | 0.82 | MMVTNNTFTL | 0.78 | 0.49 | 50 | 1 |
| S protein | NLNESLIDL | PC | 0.28 | 1.27 | NLNESLIDL | 0.29 | 1.27 | 100 | 2 |
| | FIAGLIAIV | NP | 0.30 | 0.09 | FIAGLIAIV | 0.26 | 0.09 | 100 | 2 |
| | ALNTLVKQL | PC | 0.34 | 3.64 | ALNTLVKQL | 0.66 | 3.64 | 100 | 2 |
| | KLPDDFMGCV | NP | 0.37 | 0.41 | KLPDDFTGCV | 0.49 | 0.73 | 90 | 2 |
| | RLNEVAKNL | NP | 0.43 | 0.86 | RLNEVAKNL | 0.51 | 0.86 | 100 | 2 |
| | LITGRLQSL | NP | 0.48 | 0.41 | LITGRLQSL | 0.66 | 0.41 | 100 | 2 |
| | NYNYKYRYL | NP | 0.48 | 0.09 | NYNYLYRLF | 0.51 | 0.39 | 67 | 1 |

| | | | | | | | | | |
|---|---|---|---|---|---|---|---|---|---|
| | GTGVLTPSSK | NP | 0.50 | 0.64 | GTGVLTESNK | 0.55 | 0.90 | 80 | 1 |
| | NCVADYSVLY | NP | 0.53 | 0.42 | NCVADYSVLY | 0.51 | 0. 42 | 100 | 1 |
| | RNFFSPQI | NP | 0.54 | 0.33 | RNFYEPQI | 0.61 | 1.41 | 75 | 1 |
| | VLNDILSRL | PC | 0.56 | 0.10 | VLNDILSRL | 0.58 | 0.10 | 100 | 2 |
| | NYKYRYLR | NP | 0.58 | 0.33 | NYLYRLFR | 0.52 | 0.32 | 63 | 1 |
| | GPKLSTDLI | NP | 0.60 | 0.95 | GPKKSTNLV | 0.74 | 0.47 | 67 | 1 |
| | SIVAYTMSL | PC | 0.63 | 0.08 | SIIAYTMSL | 0.72 | 0.04 | 89 | 2 |
| | QMYKTPTLK | NP | 0.65 | 0.01 | QIYKTPPIK | 0.48 | 0.16 | 67 | 1 |
| | DEIFRSDTL | NP | 0.67 | 0.10 | DKVFRSSVL | 0.52 | 0.03 | 56 | 1 |
| | IYSTGNNVF | NP | 0.68 | 0.16 | VYSTGSNVF | 0.77 | 0.14 | 78 | 1 |
| | KSIVAYTMS | NP | 0.71 | 0.96 | QSIIAYTMS | 0.73 | 9.17 | 78 | 1 |
| | KNKDGFLYV | NP | 0.72 | 0.29 | KNIDGYFKI | 0.68 | 0.95 | 44 | 1 |
| | QKSIVAYTM | NP | 0.75 | 0.47 | SQSIIAYTM | 0.74 | 0.09 | 67 | 1 |
| | IGAGICASY | NP | 0.80 | 0.08 | IGAGICASY | 0.71 | 0.08 | 100 | 1 |
| | IGAEHVDTSY | NP | 0.81 | 0.74 | IGAEHVNNSY | 0.79 | 0.49 | 80 | 1 |
| ORF3a/b | RIIMRCWLCW | NP | 0.41 | 0.35 | RIIMRLWLCW | 0.49 | 0.30 | 90 | 1 |
| | SITAQPVKI | PC | 0.42 | 4.48 | TVTLKQGEI | 0.31 | 6.95 | 22 | 2 |
| | ASLPFGWLV | NP | 0.69 | 0.22 | ASLPFGWLI | 0.54 | 0.07 | 89 | 1 |
| | STNLCTHSF | NP | 0.60 | 0.35 | NA | NA | NA | NA | 1 |
| E pr. | TLIVNSVLLF | NP | 0.38 | 0.58 | TLIVNSVLLF | 0.47 | 0.58 | 100 | 1 |
| | CNIVNVSLVK | NP | 0.42 | 0.71 | CNIVNVSLVK | 0.44 | 0.71 | 100 | 1 |
| M protein | TLACFVLAAV | NP | 0.27 | 0.15 | TLACFVLAAV | 0.28 | 0.15 | 100 | 2 |
| | RGTIVTRPL | NP | 0.53 | 0.54 | HGTILTRPL | 0.52 | 0.87 | 78 | 1 |
| | GLMWLSYFV | NP | 0.54 | 0.01 | GLMWLSYFI | 0.58 | 0.02 | 89 | 2 |
| | KEITVATSRT | NP | 0.63 | 0.69 | KEITVATSRT | 0.64 | 0.69 | 100 | 1 |
| | SQRVGTDSGF | NP | 0.64 | 0.06 | SQRVAGDSGF | 0.69 | 0.05 | 80 | 1 |
| | HLRMAGHSL | NP | 0.65 | 0.07 | HLRIAGHHL | 0.58 | 0.49 | 78 | 1,2 |
| | FAAYNRYR | NP | 0.71 | 0.70 | FAAYSRYR | 0.72 | 0.56 | 88 | 1 |
| | GHLRMAGHSL | NP | 0.73 | 0.34 | GHLRIAGHHL | 0.67 | 1.06 | 80 | 1 |
| | YRIGNYKL | NP | 0.80 | 0.36 | YRIGNYKL | 0.85 | 0.36 | 100 | 1 |
| | YYKLGASQR | NP | 0.80 | 0.26 | YYKLGASQR | 0.80 | 0.26 | 100 | 1 |
| | ATSRTLSY | NP | 0.83 | 0.75 | ATSRTLSY | 0.80 | 0.75 | 100 | 1 |
| ORF6 | WNLDVIISSI | NP | 0.44 | 1.00 | WNLDYIINLI | 0.49 | 1.08 | 70 | 1 |
| | AEILIIIMRTF | NP | 0.44 | 0.11 | AEILLIIMRTF | 0.51 | 0.07 | 91 | 1 |
| ORF9b | SQLSLSMAR | NP | 0.35 | 0.53 | NA | NA | NA | NA | 1 |
| | EELPDEFVV | NP | 0.36 | 0.55 | NA | NA | NA | NA | 1 |
| | LEARAFQST | NP | 0.38 | 0.37 | NA | NA | NA | NA | 1 |
| N protein | MEVTPSGTWL | PC | 0.38 | 0.13 | MEVTPSGTWL | 0.38 | 0.38 | 100 | 2 |
| | GMSRIGMEV | PC | 0.40 | 0.61 | GMSRIGMEV | 0.46 | 0.40 | 100 | 2 |
| | RLNQLESKV | PC | 0.42 | 2.95 | RLNQLESKM | 0.54 | 0.89 | 89 | 2 |
| | TKQYNVTQAF | NP | 0.46 | 0.01 | TKAYNVTQAF | 0.60 | 0.14 | 90 | 2 |
| | LQLPQGTTL | PC | 0.51 | 0.10 | LQLPQGTTL | 0.57 | 0.10 | 100 | 2 |
| | GETALALLLL | NP | 0.52 | 0.10 | GDAALALLLL | 0.58 | 2.73 | 80 | 1,2 |
| | EASLPYGANK | NP | 0.53 | 0.94 | EAGLPYGANK | 0.49 | 4.33 | 90 | 1 |
| | ALNTPKDHI | PC | 0.59 | 10.21 | ALNTPKDHI | 0.54 | 10.21 | 100 | 2 |
| | LLLDRLNQL | NP | 0.59 | 0.13 | LLLDRLNQL | 0.69 | 0.13 | 100 | 2 |
| | LALLLLDRL | NP | 0.62 | 0.99 | LALLLLDRL | 0.67 | 0.99 | 100 | 1,2 |
| | YGANKEGIVW | NP | 0.64 | 0.22 | YGANKDGIIW | 0.56 | 0.18 | 80 | 1 |
| | QFKDNVILL | NP | 0.67 | 0.24 | NFKDQVILL | 0.61 | 0.30 | 78 | 2 |
| | KTFPPTEPK | NP | 0.67 | 0.02 | KTFPPTEPK | 0.65 | 0.02 | 100 | 1 |
| | SPRWYFYYLG | NP | 0.68 | 0.40 | SPRWYFYYLG | 0.59 | 0.40 | 100 | 1 |
| | DAYKTFPPT | NP | 0.70 | 0.74 | DAYKTFPPT | 0.66 | 0.74 | 100 | 1 |

[1]MAPs are classified as either naturally processed (NP) or potential candidates (PC).

**Supplementary Table S3. SARS-CoV-2 specific MAPs identified with mass spectrometry analyses of SARS-CoV-2 infected cells (**Weingarten-Gabbay, S. et al 2020) **that were in our target list.**

| Protein | Mass spectrometry | | | Target list | | |
|---|---|---|---|---|---|---|
| | Peptide sequence | CAMAP score | Rank | Peptide sequence | CAMAP score | Rank |
| S | VG**YLQPRTF** | 0.543 | 0.235 | **YLQPRTF**LL | 0.68 | 0.017 |
| M | **VATSRTL**SY | 0.667 | 0.086 | IT**VATSRTL** | 0.674 | 0.134 |
| ORF1ab | **STSAFV**ETV | 0.842 | 0.035 | FSA**STSAFV** | 0.787 | 0.01 |

**Supplementary Table S4. List of top SARS-CoV-2 target peptides as predicted using only the cumulative frequency of haplotypes.** Only 10/48 (29.2%) of MAPs were homologous (or with an overlap >50%) to those that were previously identified as naturally processed SARS-CoV-1 MAPs[1,28] (indicated with a star: *). Four MAPs (shown in bold) were also identified when filtering for high CAMAP scores (i.e. above the median CAMAP score of its gene of origin, see Table 1).

| Protein | Peptide sequence | CAMAP score | Min rank (%) | #alleles[1] | #haplotypes[2] | Haplotypes freq. (%) in population[3] |
|---|---|---|---|---|---|---|
| Orf1ab (Replicase) | | 0.560 | 0.022 | 9 | 4176 | 57.8% |
| | TLMNVLTLVY* | 0.701 | 0.068 | 4 | 2886 | 54.1% |
| | YMPYFFTLL | 0.667 | 0.010 | 8 | 3844 | 53.4% |
| | FSASTSAFV | 0.787 | 0.010 | 7 | 3539 | 50.6% |
| | FVDGVPFVV* | 0.543 | 0.016 | 7 | 3539 | 50.6% |
| | FLYENAFLPF | 0.685 | 0.021 | 8 | 4049 | 50.0% |
| | MSAFAMMFV | 0.765 | 0.007 | 6 | 3128 | 48.2% |
| | FLGRYMSAL | 0.693 | 0.099 | 7 | 3248 | 46.7% |
| | FLLNKEMYL | 0.534 | 0.008 | 7 | 3248 | 46.7% |
| | FLNRFTTTL | 0.472 | 0.010 | 7 | 3248 | 46.7% |
| S protein | MIAQYTSAL | 0.596 | 0.015 | 9 | 3904 | 59.4% |
| | **SIIAYTMSL**** | 0.718 | 0.039 | 8 | 3535 | 54.7% |
| | YLQPRTFLL | 0.680 | 0.017 | 8 | 3844 | 53.4% |
| | STQDLFLPF | 0.542 | 0.118 | 8 | 3679 | 45.0% |
| | FQFCNDPFL | 0.575 | 0.036 | 6 | 3109 | 38.6% |
| | FTISVTTEI | 0.429 | 0.009 | 6 | 3109 | 38.6% |
| | VVFLHVTYV | 0.382 | 0.211 | 6 | 3109 | 38.6% |
| | FTNVYADSF | 0.523 | 0.071 | 7 | 3108 | 38.4% |
| | YSSANNCTF | 0.714 | 0.013 | 7 | 3108 | 38.4% |
| | KIADYNYKL | 0.447 | 0.202 | 5 | 2569 | 34.2% |
| M protein | RLFARTRSM | 0.476 | 0.038 | 6 | 2561 | 39.6% |
| | YANRNRFLY | 0.273 | 0.027 | 6 | 2605 | 34.8% |
| | **ATSRTLSYY*** | 0.807 | 0.008 | 4 | 2175 | 33.5% |
| | FLWLLWPVTL | 0.420 | 0.211 | 2 | 2028 | 33.2% |
| | FVLAAVYRI | 0.300 | 0.125 | 3 | 1815 | 28.0% |
| | AIAMACLVGL | 0.211 | 0.852 | 1 | 1157 | 24.1% |
| | FIASFRLFA | 0.479 | 0.232 | 1 | 1157 | 24.1% |
| | FLFLTWICLL | 0.213 | 0.348 | 1 | 1157 | 24.1% |
| | FLYIIKLIFL | 0.252 | 0.814 | 1 | 1157 | 24.1% |
| | GLMWLSYFI* | 0.580 | 0.021 | 1 | 1157 | 24.1% |
| N protein | FPRGQGVPI | 0.538 | 0.009 | 5 | 2089 | 36.7% |
| | FGMSRIGMEV* | 0.474 | 0.325 | 4 | 1989 | 36.4% |
| | **LLLDRLNQL*** | 0.694 | 0.134 | 3 | 1759 | 34.9% |
| | FAPSASAFF | 0.339 | 0.016 | 7 | 3266 | 33.2% |
| | KAYNVTQAF* | 0.562 | 0.008 | 6 | 2695 | 26.5% |
| | SPRWYFYYL* | 0.483 | 0.009 | 3 | 1041 | 23.6% |
| | LLNKHIDAY | 0.476 | 0.101 | 3 | 1538 | 22.3% |
| | LTYTGAIKL | 0.421 | 0.128 | 5 | 2281 | 21.0% |
| | NTASWFTAL | 0.477 | 0.032 | 5 | 2281 | 21.0% |
| | KPRQKRTAT | 0.539 | 0.065 | 2 | 522 | 19.9% |
| E protein | YVYSRVKNL | 0.502 | 0.037 | 6 | 2438 | 30.0% |
| | FLLVTLAIL | 0.478 | 0.347 | 3 | 1868 | 29.4% |
| | FLAFVVFLL | 0.469 | 0.061 | 3 | 1815 | 28.0% |
| | FVSEETGTL | 0.537 | 0.028 | 6 | 2695 | 26.5% |
| | SLVKPSFYV | 0.471 | 0.097 | 1 | 1157 | 24.1% |
| | SVLLFLAFV | 0.406 | 0.413 | 1 | 1157 | 24.1% |
| | **IVNSVLLFL*** | 0.424 | 0.230 | 5 | 2281 | 21.0% |
| | LAILTALRL | 0.611 | 0.117 | 5 | 2281 | 21.0% |

**Supplementary Table S5. HLA allele restriction for each of the top SARS-CoV-2 target peptides.**

| Prot. | Peptide | Allele restriction[1] | | |
|---|---|---|---|---|
| | | HLA-A | HLA-B | HLA-C |
| Orf1ab (Replicase) | FSASTSAFV | 01:01, 02:01 | | 01:02, 02:02, 03:04, 08:01, 16:01 |
| | MSAFAMMFV | 01:01, 02:01 | | 02:02, 03:04, 08:01, 16:01 |
| | YLITPVHVM | 02:01 | 35:01 | 01:02, 02:02, 03:04, 08:01, 16:01 |
| | MVMCGGSLY | 01:01, 03:01, 11:01 | 35:01 | 02:02, 16:01 |
| | FLTENLLLYI | 01:01, 02:01 | | 02:02 |
| | FSSTFNVPM | 01:01 | 35:01 | 01:02, 02:02, 03:04, 08:01, 16:01 |
| | MADQAMTQM | 01:01 | 35:01 | 01:02, 02:02, 03:04, 08:01, 16:01 |
| | KLVSSFLEM | 02:01 | | 01:02, 02:02, 03:04, 16:01 |
| | AVIKTLQPV | 02:01 | | 02:02, 03:04, 08:01, 16:01 |
| | FISTCACEI | 02:01 | | 01:02, 02:02, 03:04, 08:01 |
| S protein | SIIAYTMSL | 02:01 | 07:02, 08:01 | 01:02, 02:02, 03:04, 08:01, 16:01 |
| | YLQPRTFLL | 02:01, 24:02 | 08:01 | 01:02, 02:02, 03:04, 08:01, 16:01 |
| | YSSANNCTF | 01:01 | 35:01 | 01:02, 02:02, 03:04, 08:01, 16:01 |
| | AALQIPFAM | | 35:01 | 01:02, 02:02, 03:04, 08:01, 16:01 |
| | RLDKVEAEV | 02:01 | | |
| | RLQSLQTYV | 02:01 | | |
| | TSNQVAVLY | 01:01 | 35:01 | 02:02 |
| | CSFGGVSVI | | | 01:02, 02:02, 03:04, 08:01, 16:01 |
| | IANQFNSAI | | | 01:02, 02:02, 03:04, 08:01, 16:01 |
| | TSVDCTMYI | | | 01:02, 02:02, 03:04, 08:01, 16:01 |
| M protein | ATSRTLSYY | 01:01, 03:01, 11:01 | | 02:02 |
| | FAAYSRYRI | | | 01:02, 02:02, 03:04, 08:01, 16:01 |
| | ITVATSRTL | | | 01:02, 02:02, 03:04, 08:01, 16:01 |
| | VAGDSGFAAY | 01:01 | 35:01 | |
| | RIAGHHLGR | 03:01, 11:01 | | |
| | YSRYRIGNYK | 03:01, 11:01 | | |
| | WSFNPETNI | | | 02:02, 03:04, 08:01 |
| | ESELVIGAVI | | 44:02, 44:03 | |
| | RPLLESELVI | | 07:02 | |
| | VPLHGTIL | | 07:02 | |
| N protein | LLLDRLNQL | 02:01 | 08:01 | 01:02 |
| | KTFPPTEPKK | 03:01, 11:01 | | |
| | LPNNTASWF | | 07:02, 35:01 | |
| | FSKQLQQSM | | | 02:02, 03:04, 16:01 |
| | LQLPQGTTL | | | 03:04, 08:01, 16:01 |
| | LSPRWYFYY | 01:01 | | |
| | QGTTLPKGFY | 01:01 | | |
| | SSPDDQIGYY | 01:01 | | |
| | RIRGGDGKMK | 03:01 | | |
| | KFPRGQGVPI | | 07:02 | |
| E protein | YVYSRVKNL | | 08:01 | 01:02, 02:02, 03:04, 08:01, 16:01 |
| | FLLVTLAIL | 02:01 | | 03:04, 08:01 |
| | FLAFVVFLL | 02:01 | | 02:02, 08:01 |
| | FVSEETGTL | | 35:01 | 01:02, 02:02, 03:04, 08:01, 16:01 |
| | LAILTALRL | | | 01:02, 02:02, 03:04, 08:01, 16:01 |
| | SLVKPSFYVY | 03:01 | | |
| | LIVNSVLLF | | 35:01 | 02:02 |
| | FVVFLLVTL | | | 03:04 |
